# Stress-responsive roles of the *C. neoformans* human-like eIF3 complex

**DOI:** 10.64898/2026.01.12.698982

**Authors:** Maritza N. Ventura, Prabhakar Singh, David Goich, John C. Panepinto

## Abstract

Eukaryotic translation initiation factor 3 (eIF3) is a complex of proteins that plays a pleiotropic role in translation regulation across eukaryotes, but the composition of eIF3 complexes varies with retention and loss of subunit genes across evolution. The model yeast *Saccharomyces cerevisiae* encodes six eIF3 subunits whereas mammals encode thirteen subunits. The basidiomycete fungus and opportunistic fungal pathogen, *Cryptococcus neoformans*, encodes a mammalian complement of eIF3 subunits. In this report, we investigated the contribution of the non-essential eIF3 subunit genes to cryptococcal stress tolerance. We found that mutants in the four nonessential subunits, eIF3d, eIF3e, eIF3k and eIF3l all exhibit sensitivity to mitochondrial perturbation, and that mutants in eIF3d and eIF3e exhibit opposite susceptibilities to the antifungal drug fluconazole and the hypoxia mimetic cobalt chloride. Loss of eIF3d resulted in reduced eIF2α phosphorylation in response to stress, but the mutant was still able to repress translation to the same extent as the wild type and was defective in induction of integrated stress response regulon. Despite producing higher levels of urease and melanin, the eIF3d deletion mutant was avirulent in *Galleria mellonella* larvae. Together our data demonstrates the importance of *C. neoformans* eIF3 in stress adaptation and pathogenesis.

**Importance:** Cryptococcus *neoformans* is an opportunistic fungal pathogen that causes cryptococcal meningoencephalitis in immunocompromised individuals leading to ∼120,000 deaths worldwide annually. When *C. neoformans* is exposed to host-relevant stressors, such as oxidative stress and thermal stress, it reprograms the translating pool of mRNAs to favor stress adaptation. Eukaryotic translation initiation factor 3 is a multi-subunit complex with roles in stress-responsive translation across eukaryotes yet is unexplored in *C. neoformans.* We found that *C. neoformans* encodes orthologues of all thirteen mammalian eIF3 subunits. Mutational analysis of non-essential subunits implicated eIF3 in responses to mitochondrial stress and antifungal susceptibility in *C. neoformans,* and demonstrates a role for eIF3d in the induction of the integrated stress response as well as in *Cryptococcal* pathogenesis. Further work will investigate the specific mRNAs that are regulated by eIF3 in response to host-relevant stressors in *C. neoformans*.

## Introduction

Environmental fungi with human pathogenic potential exhibit high degrees of adaptability. The transition from the soil environment to the human host is associated with thermal stress resulting from mammalian endothermy, nutritional immunity that requires metabolic adaptation, and oxidative and nitrosative insults from innate immune cells (1). Investigation into the molecular mechanisms governing stress adaptation in fungi have focused on transcriptional regulation, revealing intricate networks of transcription factors that orchestrate gene expression in response to various stimuli (2). Less understood are the mechanisms of translational regulation that impact stress adaptation in fungi.

Our work has focused on post-transcriptional processes that promote stress adaptation in the human fungal pathogen *C. neoformans,* which is the cause of approximately 19% of HIV-associated mortality and remains a significant cause of invasive infection in people living with solid organ transplants (3). Our work has led us to a model of translatome reprogramming in which mRNA decay and regulation of translation initiation act in parallel to promote the prioritization of stress responsive mRNAs (4–6). Additional translational regulatory mechanisms likely impact stress adaptation in *C. neoformans* and other fungal pathogens (7–9).

In this study we focus on eukaryotic initiation factor 3 (eIF3), which is a multi-subunit initiation factor that plays a variety of roles in translation across eukaryotic cells (5). Interestingly, the subunit composition of eIF3 varies across evolution, with the model yeast *S. cerevisiae* encoding six subunits (10, 11), and mammals encoding thirteen subunits (12). Other model fungi encode a more mammalian complement of subunits, with the fission yeast *Schizosaccharomyces pombe* encoding eleven subunits (13), and the model mold *Neurospora crassa* encoding all twelve (14). The function of eIF3 in human fungal pathogens has not been explored.

eIF3 influences multiple steps in the initiation of translation, interacting with the cap to promote mRNA recruitment and bridging the 40S ribosomal subunit to stabilize the initiation complex (10). The core of mammalian eIF3 is comprised of subunits a, c, e, l, k, and m, all of which contain a PCL domain that promotes core formation, and two MPN-domain containing proteins, eIF3f and h, that associate via interactions with eIF3m (11, 15). Although the *S. cerevisiae* eIF3 lacks the PCL domain proteins eIF3e, l and k, *S. cerevisiae* eIF3 a and c are sufficient to form the core complex that associates with the mRNA exit channel of the ribosome, and can wrap around the ribosome via eIF3b to reach the mRNA entry channel (16). The major differences, therefore, between the mammalian and *S. cerevisiae* eIF3 are in the structural complexity of the PCL core, and in the association of eIF3h, f and d with the PCL core on the mRNA exit side of the 40S ribosome subunit (10).

In this study, we assess the eIF3 diversity across fungal pathogens, and find variations in subunit components across the fungal pathogens with the basidiomycete yeast *C. neoformans* containing the full mammalian complement of eIF3 subunits. To investigate the function of eIF3 in *C. neoformans*, we screened available deletion mutants in eIF3d, e, k, and l for stress sensitivity and thermotolerance. We discovered variable stress phenotypes in the four subunit mutants, with loss of eIF3d conferring the most drastic stress-sensitivity. Further investigation of translational stress response in the *eIF3d*Δ mutant revealed normal translational repression in response to stress, but a defect in activation of Gcn2, leading to a defect in induction of the integrated stress response. Although the *C. neoformans eIF3d*Δ mutant produced increased amounts of the virulence factors urease and melanin, the mutant was avirulent in the *G. mellonella* model of infection.

## Materials and Methods

### Evolutionary Analysis

To identify subunits of eIF3 present in human pathogenic fungi and model fungi, we performed a protein BLAST search of eIF3 subunits in *C. neoformans* and *N. crassa* (14). These protein sequences were obtained via FungiDB (release 68) for *N. crassa* OR74A and *C. neoformans* H99, with the exception of eIF3b/PRT1 in *C. neoformans*, which was found under the GenBank identifier AAN75151.2. The BLASTp suite was accessed via the NCBI BLAST portal. The search set was specified against the non-redundant protein set (nr) and limited to the following species/IDs: *C. neoformans* (taxid: 235443), *Cryptococcus amylolentus* (taxid:104669), *Coprinopsis cinereus* (taxid:5346), *Mycosarcoma* (or *Ustilago*) *maydis* (taxid:5270), *Aspergillus fumigatus* (taxid:746128), *Candidozyma auris* (taxid:498019), *Candia albicans* (taxid:5476), *Nakaseomyces glabratus* (*Candida glabrata*) (taxid:5478), *S. cerevisiae* (taxid:4932), *S. pombe* (taxid:4896), *N. crassa* (taxid:5141). The BLAST search was specified to return 500 target sequences and was otherwise run with default parameters. As a complementary approach, we cross-referenced our results with gene annotations for these species in FungiDB, using orthology tables to identify subunit homologs which were missed by our BLAST search. The taxonomic relation between these fungi was obtained using the NCBI Taxonomy Common Tree tool and visualized alongside our BLAST results in R. Clustal Omega alignments of protein sequences were used to create percent identity matrices and phylogenetic trees through the UniProt interface. Protein sequence identifiers are provided in Table S1.

### Strains and Media

*C. neoformans* strain KN99, the parental for the Madhani Deletion Collection, and the collection of deletion mutants in this genetic background were purchased from the Fungal Genetics Stock Center at the University of Kansas. After initial phenotyping, the collection *eIF3d*Δ mutant wascomplemented by reintroduction of the wild type gene in trans. The coding region and 1kb upstream and down-stream was amplified by PCR using primers eIF3d-Comp-SpeI-Fwd (5’-AAAAAAACTAGTCCTATAGAAGAAAGTTGGG-3’) and eIF3d-Comp-SpeI-Rev (5’- AAAAAAACTAGTGGTAAATACTCACGACAGATGG-3’), digested with *Spe* I and ligated into *Spe* I-digested alkaline phosphatase-treated pBluescript that also contained the G418 resistance cassette.

### Northern Blot Analysis

Midlogarithmic phase (OD_600_ = 0.6) grown cells (15 mL) of selected *C. neoformans* strains with respective treatment and culture conditions were pelleted by centrifugation at 4000 rpm for 2 minutes, 50 µl RLT buffer with 2-mercaptoethanol (10 µl/mL) was added to pellets and were flash frozen in liquid nitrogen. RNA extraction and northern blotting were performed as described by C. M. Knowles et al. (4). In brief the pellets were thawed and lysates were sandwiched between glass beads in centrifuge tubes. The pellets were lysed mechanistically using a Bullet Blender and lysate removed from the bead bed by centrifugation. Whole-cell lysates were cleared by centrifugation at 15000 rpm for 10 min and the supernatants were transferred to fresh microcentrifuge tubes where equal volumes of 70% ethanol were added to precipitate the nucleic acids. Qiagen RNeasy kit was used to extract RNA according to the manufacturer’s instructions. RNA was quantified by NanoDrop, and 3 µg was loaded onto a formaldehyde-agarose gel for electrophoresis. RNA was visualized on a Bio-Rad Gel Doc and and rRNA was quantified using ImageLab software. RNA was transferred to a Hybond-nylon membrane and probed using an α-^32^P-labeled DNA probe, created using Invitrogen RadPrime DNA Labeling kit (cat no. 18428011). The *LEU2* probe was generated using genomic DNA amplified using LEU2 Northern forward (CGAGACTATTGTCAACCATTCTG) and reverse (CTTCTTCACAGCATGCTC) primer as described by Panepinto, et al., (2002). Typhoon phosphoimager was used for phosphor scanning and imaging of the blot, using a GE Typhoon phosphoimager and hybridized signal was quantified using ImageLab software.

### Polysome Profiling

Polysome profiling was performed as described in detail by C. M. Knowles et al. (17). Briefly, overnight-grown wt and mutant strains of *C. neoformans* were inoculated into fresh YPD medium at an initial OD_600_ of 0.2 and grown to the mid-logarithmic phase under optimal growth conditions. Control samples from both wt and mutant strains were collected, and the remaining cells were subjected to oxidative stress (1 and 1.5 mM H₂O₂ for 30 min) or thermal stress (37°C for 30 and 60 min). Cells were pelleted, flash-frozen, and resuspended in 1 mL of lysis buffer (20 mM Tris-HCl, pH 8.0; 140 mM KCl; 5 mM MgCl₂; 1% Triton X-100; 0.5 mg/mL heparin sodium sulfate; and 0.1 mg/mL cycloheximide) and transferred to microcentrifuge tubes. Samples were centrifuged at 4,000 rpm for 3 min, the pellets were resuspended in 50 µL of polysome lysis buffer, and glass beads were added for cell lysis using a pre-cooled Bullet Blender with dry ice (5 min at setting 12). An additional 150 µL of lysis buffer was added, and the lysate was centrifuged at 15,000 rpm for 10 min. The clarified lysate was collected, and RNA concentration was determined using a NanoDrop spectrophotometer (Thermo Fisher Scientific). For each profile, 150 µg of RNA was layered onto a 10–40% sucrose gradient and centrifuged at 39,000 rpm for 120 min in an SW41Ti swinging-bucket rotor. RNA absorbance at 260 nm was recorded using Biocomp gradient fractionator coupled with Triax flow cell. Graphs were generated using GraphPad Prism version 10.6.1.

### Spot Plate Assay

The spot plate assay was performed following the methodology described by A. L. Bloom et al. (18). Briefly, the KN99, *eIF3dΔ* and *eIF3dΔ*::eIF3D strains were seeded in YPD broth and incubated at 30°C overnight. Overnight grown cells were harvested by centrifugation at 4000 rpm for 2 min, washed with sterile deionized water (SDW), then resuspended in SDW at OD_600_ = 1.0. A ten-fold serial dilution was prepared for up to five dilution steps (10^−5^), and 5 μl of each dilution was spotted on the YPD agar plates containing indicated doses of antifungal agents. The plates were incubated at 30°C and/or 37°C for 72 hours and photographed.

### Western blotting

Midlogarithmic phase (OD_600_ = 0.6) grown cells (15 mL) of selected *C. neoformans* strains with respective treatment and culture conditions were pelleted by centrifugation at 4000 rpm for 2 min, and cell pellets were flash frozen in liquid nitrogen. Protein extraction and SDS-PAGE were performed as described by C. M. Knowles et al. (4). To detect phosphorylated eIF2α, blots were incubated with rabbit anti-phospho-eIF2α (Abcam, 1:1000), followed by anti-rabbit Horseradish peroxidase (HRP) conjugated secondary antibody (1:10000) (Cell Signaling). Blots were stripped and incubated with rabbit anti-eIF2α (Genscript) or RPS6 (Abcam, 1:1000), followed by HRP-conjugated anti-rabbit secondary Ab. For Gcn4, blot was probed using a polyclonal rabbit anti-Gcn4 antibody (1:1000, Genscript) followed by HRP-conjugated anti-rabbit secondary Ab. Blots were imaged on a Bio Rad Chemidoc and chemiluminescent signals were quantified using ImageLab software (Bio-Rad).

### Virulence factors (urease and melanin)

To analyze melanin and urease production, overnight grown cells were washed twice with SDW and resuspended in 10 mL SDW to OD_600_ = 1.0. Cells were pelleted at 4000 rpm for 2 min, residual water was aspirated, and cells were resuspended in 100 µL SDW. For urease assessment 10 µL cells from each strain were spotted on urea base agar (peptone (1g/L), glucose (1g/L), sodium chloride (5g/L), Di-sodium phosphate (1.2g/L), potassium dihydrogen phosphate (0.8g/L), phenol red (0.012g/L), agar (15g/L)) plates and incubated at 30°C and 37°C for overnight. For laccase activity, 10 µL cells from each strain were spotted on Asparagine agar (K_2_HPO_4_ (0.5 g/L), asparagine (0.5 g/L), agar (15 g/L), pH 7.0) plate supplemented with 1 mM L-Dopamine and 0, 0.1 and 2% dextrose. Plates were incubated at 30°C and 37°C overnight and photographed after 24 hrs.

## *G. mellonella* Survival Assay

Overnight grown cultures of *C. neoformans* strains (H99, *eIF3dΔ* and *eIF3dΔ*::eIF3D) were washed twice with phosphate-buffered saline (PBS) and resuspended in 1 mL PBS. Cells were counted using a hemocytometer and the inoculum was prepared by diluting the cells to 10^4^ cells/µl. For each strain 12 *G. mellonella* larvae (150-200 mg) were infected by injecting 10^5^ cells (10 µl) into the last left proleg. The larvae were incubated at 37°C for seven days. For control 10 µl PBS was injected in the larvae. The experiment was performed in biological triplicate. Survival curves were plotted using GraphPad Prism 10 software and Kruskal-Wallis test (ANOVA on ranks) was used to compare median survival among groups, and Dunn’s test was used in pairwise analyses post hoc. A P-value less than 0.05 was considered significant.

## Results

### *C. neoformans* encodes a mammalian eIF3 complex

We assessed the *C. neoformans* genome for the presence of eIF3 subunits and found that 12 of the 13 subunits present in the mammalian eIF3 complex were found in *C. neoformans*. Interestingly, the one subunit that lacks an annotated locus in *C. neoformans, PRT1* or eIF3b, was found by us and others to be present in the mating locus, but has not been assigned a locus identifier (19, 20). The model yeast *S. cerevisiae* is reported to contain only six eIF3 subunits in its genome, the core subunits of eIF3a, eIF3b and eIF3c, as well as eIF3g, eIF3i and eIF3j (11, 15). We set out to understand the complexity of eiF3 subunits across the model and pathogenic fungi, and found that *Nakeseomyces glabratus* (formerly *C. glabrata*) genome contains the same complement of eIF3 subunits as *S. cerevisiae*. *C. albicans* and *C. auris*, contain the six subunits of *S. cerevisiae* with the addition of eIF3f, eIF3h and eIF3m. *S. pombe* also expresses eIF3f, eIF3h and eIF3m, and also expresses eIF3d and eIF3e, but lacks eIF3k and eIF3l of the mammalian complex. Interestingly, the filamentous ascomycetes *A. fumigatus* and *N. crassa* contain the full mammalian complement, which for *N. crassa* has been previously reported (14). Similarly, the basidiomycetes investigated, including *C. neoformans* and its thermosensitive environmental relative *C. amylolentus,* as well as the corn smut pathogen *Ustilago maydis* and the model mushroom *Coprinopsis cinerea* all contain the full mammalian eIF3 complement in their genomes (Fig. 1A).

**Figure 1:**
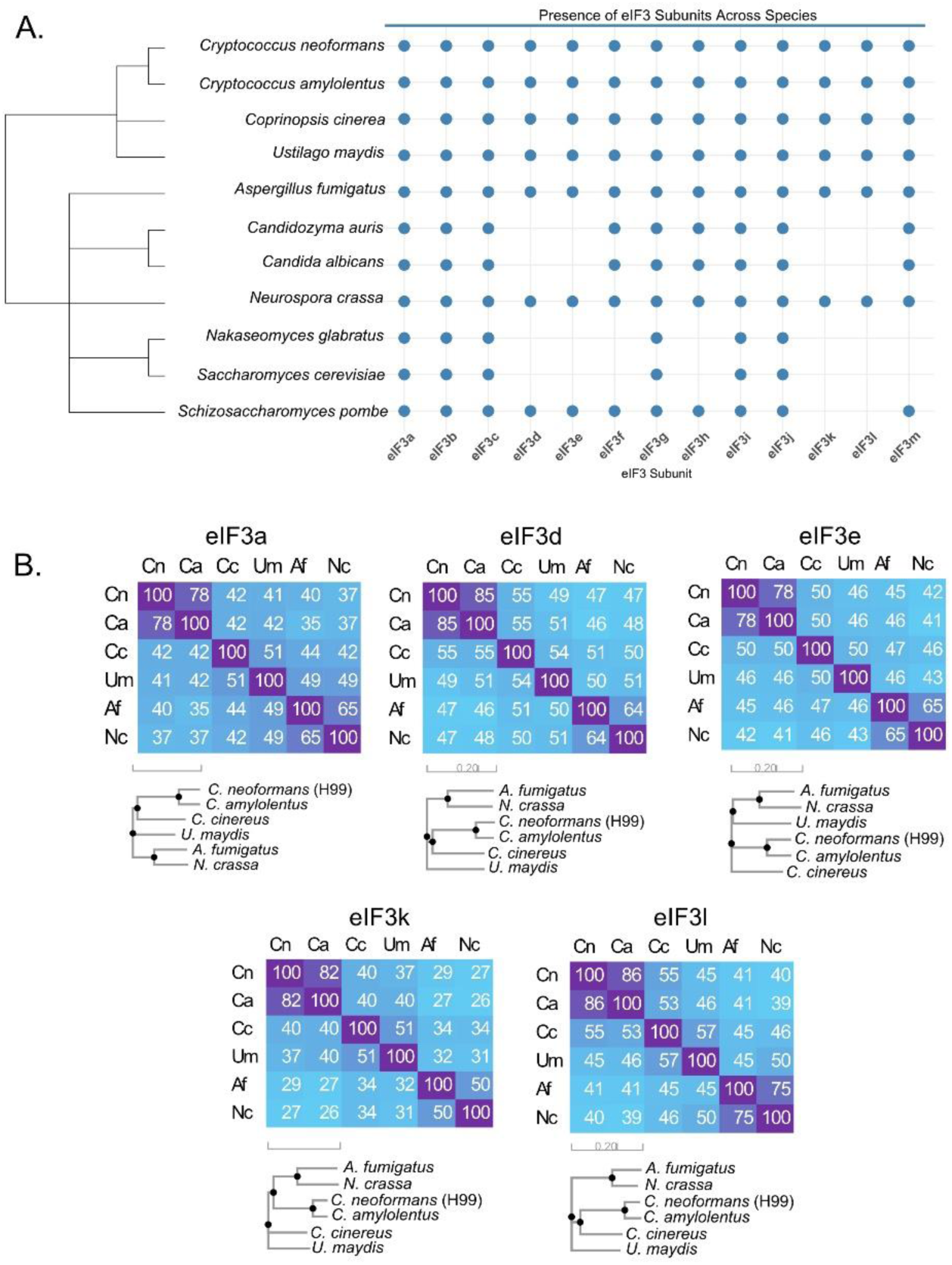
Basidiomycetes and filamentous ascomycetes encode the mammalian complement of eIF3 subunits. (A) Chart depicting the evolutionary distribution of subunits of eIF3 across the human pathogenic and model fungi. Cladogram on the left represents taxonomic relationships between species. (B) Amino acid identity matrices and homology trees for the core subunit eIF3a, and the four of the non-core subunits that are retained in mammals, basidiomycetes and filamentous ascomycetes.

We then investigated protein identity across fungi for the core subunit eIF3a, and the four subunits present in the filamentous ascomycetes and basidiomycetes (Fig. 1B). Interestingly, eIF3d and eIF3l exhibited the most protein identity across species, with 85% and 86% identity between *C. neoformans* and *C. amylolentus,* respectively, and a lowest identity of 47 % and 40% between *C. neoformans* and *N. crassa.* Although eIF3k exhibited high identity between *C. neoformans* and *C. amylolentus (82%)*, it exhibited only 50% identity in the filamentous ascomycetes (*A. fumigatus* vs. *N. crassa*) and only 27% identity between *C. neoformans* and *N. crassa*. Despite absolute conservation of eIF3a across all eukaryotes, it did not exhibit higher identity across fungi than the non-conserved subunits.

### Nonessential eIF3 subunits contribute differentially to stress adaptation in *C. neoformans*

The core subunits of eIF3 are essential for growth across eukaryotes (16, 21, 22). For the sake of clarity, we will refer to the *C. neoformans* eIF3 components by their subunit designations. Corresponding gene names based on other fungi can be found in Table S2. To investigate the function of subunits outside of the core complex, we turned to the *C. neoformans* deletion collection. Deletion mutants corresponding to four eIF3 subunits, eiF3d, eIF3e, eIF3k and eIF3l, were available. We assessed this panel of eIF3 mutants for sensitivity to various stressors and found differential sensitivity between the mutants. In response to mitochondrial stress inducers, the *eIF3d*Δ mutant exhibited sensitivity to both the redox inhibitor auranofin and hypoxia mimetic cobalt chloride at 30°C that was exacerbated by incubation at 37°C, although the *eIF3d*Δ mutant exhibited thermosensitivity at 37°C on YPD in the absence of additional stressors (Fig. 2A). At 37°C the *eIF3kΔ* mutant exhibited sensitivity to auranofin, whereas *eIF3eΔ*, *eIF3kΔ* and *eIF3IΔ* mutants showed modest resistant to cobalt chloride. All four mutants exhibited a strong sensitivity to antimycin A at 37°C with little impact at 30°C. Finally, *eIF3d*Δ mutant alone exhibited a high degree of sensitivity to the alternative oxidase inhibitor SHAM, but only at 37°C. This suggests that there may be perturbations in mitochondrial adaptation in the absence of eIF3 components. In response to peroxide, only the *eIF3dΔ* mutant exhibited sensitivity by reduced colony size, that was more pronounced at 37°C (Fig. 2B), as well as sensitivity to the antifungal drugs fluconazole and caspofungin (Fig. 2C).

**Figure 2.**
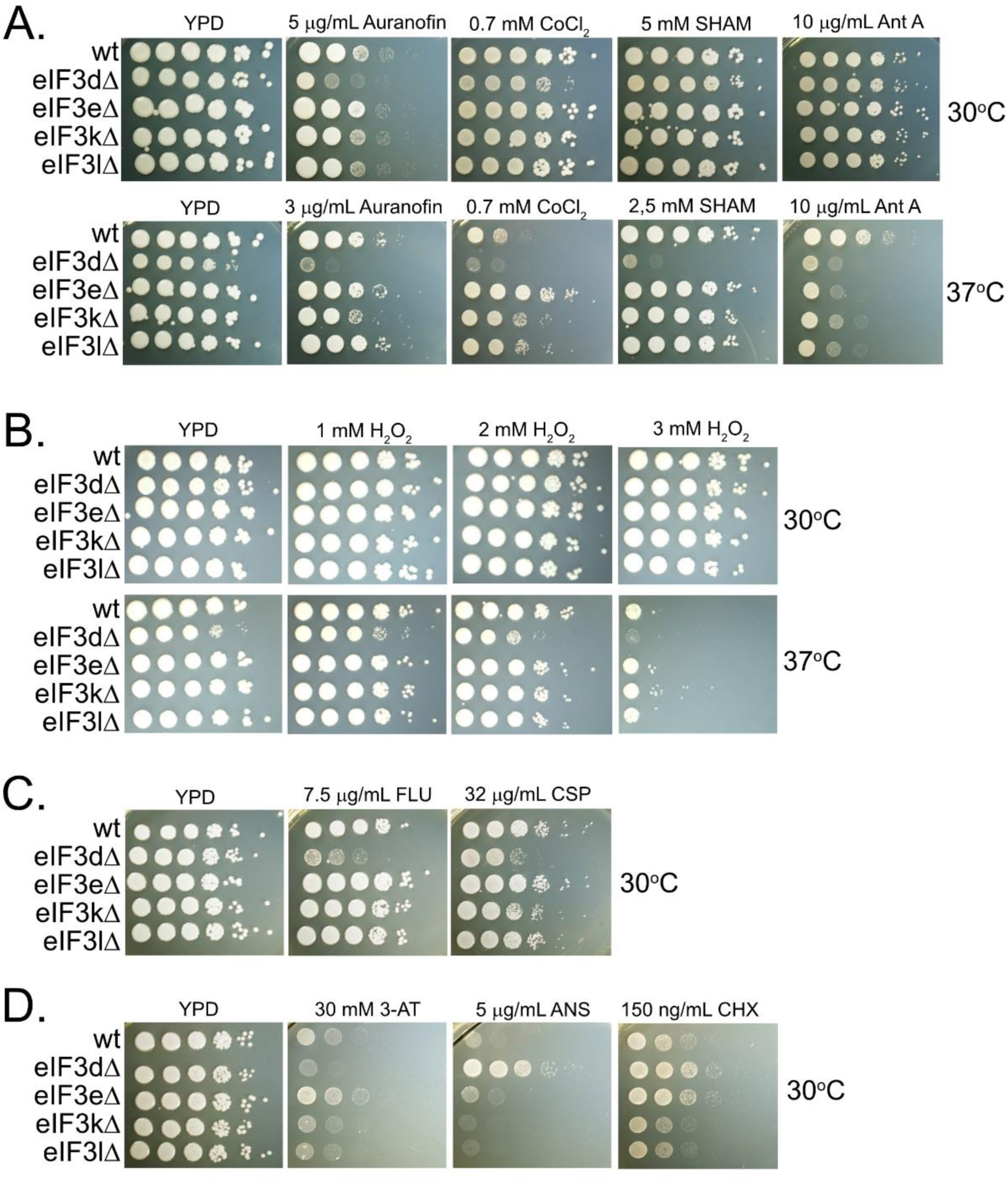
*C. neoformans* mutants of non-core eIF3 subunits vary in stress sensitivity. Spot dilution assays of wild type (KN99) and deletion mutants of eIF3d, eIF3e, eIF3k, and eIF3l on (A) mitochondrial stressors at 30°C (top) and 37°C (bottom), (B) oxidative stress (H_2_O_2_) at 30°C (top) and 37°C (bottom), (C) antifungal drugs fluconazole and capsofungin and (D) translational inhibitors. Plates were photographed after 3 days of incubation at the indicated temperature. Images shown are representative of three biological replicates.

### Deletion of eIF3d alters Gcn2 activation

Given the known role of eIF3 in translation regulation, we assessed the panel of deletion mutants for sensitivity to known translation inhibitors, and found that *eIF3d*Δ, *eIF3k*Δ and *eIF3l*Δ mutants were all sensitive to the histidine analog 3-amino triazole (3-AT) that is a known activator of the Gcn2 pathway that responds to uncharged tRNAs and ribosome collisions (Fig. 2D). Only *eIF3d*Δ mutant exhibited resistance to anisomycin, which suggests a defect in termination resulting in ribosome readthrough. In addition, *eIF3d*Δ and *eIF3e*Δ mutants exhibited resistance to cycloheximide, also suggestive of inappropriate ribosome processivity and termination defects.

Deletion of eIF3d results in both thermosensitivity and sensitivity to oxidative stress, both of which are relevant to infection, and so we set out to characterize the role of eIF3d in the translational response of *C. neoformans* to these stressors. Our first step in investigating the function of eIF3d in *C. neoformans* stress adaptation was to generate a complemented strain to assure that stress-sensitive traits were a result of the intended mutation. Re-introduction of wild type eIF3d as a transgene linked to the G418-resistance cassette restored wild type growth under oxidative stress and thermal stress (Fig. 3A). Our previous work demonstrates an alteration in the translational state in response to thermal and oxidative stress, with a more severe translational repression and robust Gcn2 activation occurring in response to oxidative stress (4). We used polysome profiling to visualize the translational state of wild type and *eIF3d*Δ in response to both thermal and oxidative stress (Fig. 3B). Under unstressed conditions, the translational state of the *eIF3d*Δ mutant is indistinguishable from wild type, and the translational repression associated with both thermal stress and oxidative stress was intact in the absence of eIF3d.

**Figure 3.**
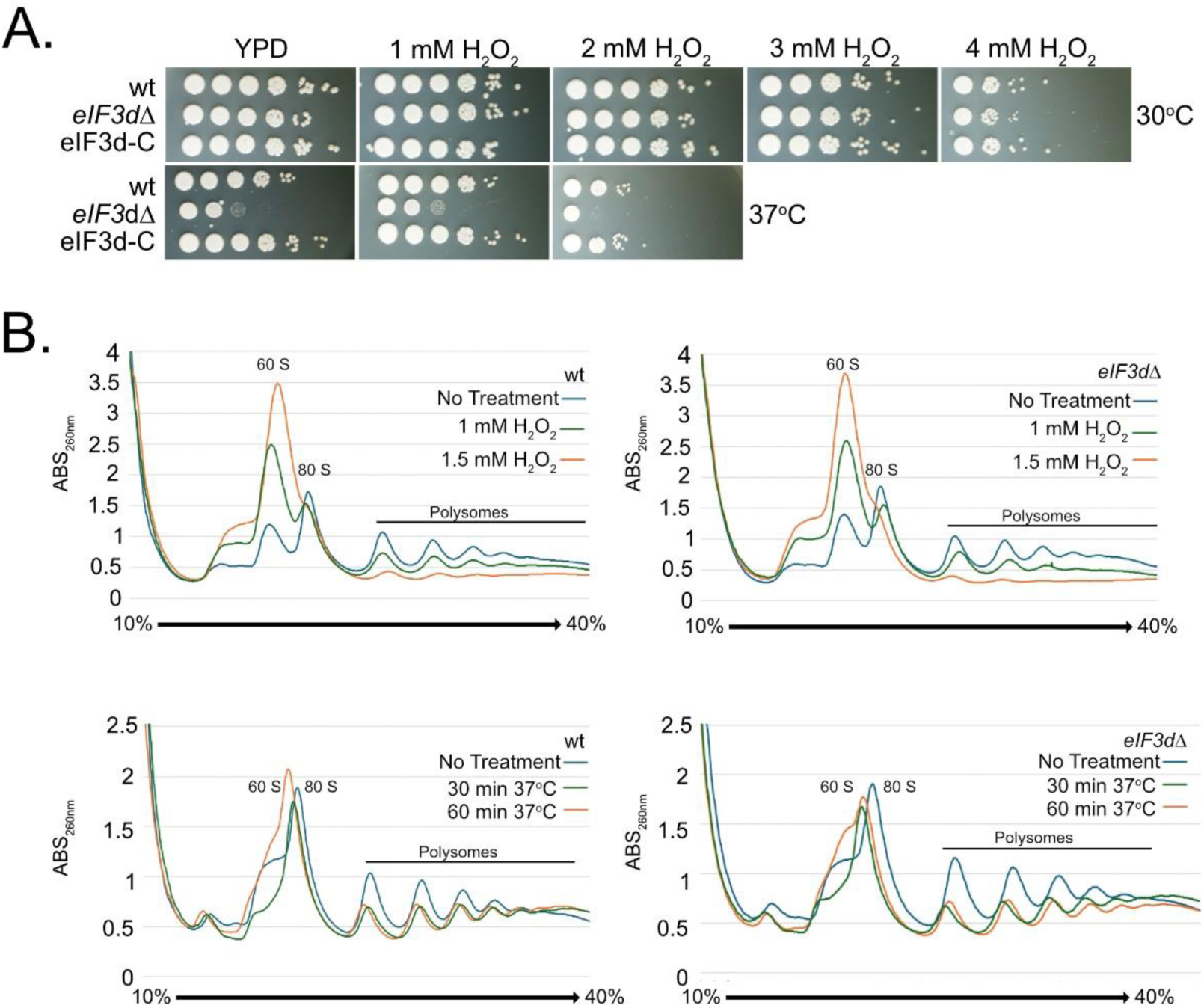
eIF3d exhibits a wild type translational response to acute oxidative and thermal stress. (A) Spot dilution of wild type, *eIF3d*Δ and the *eIF3d* complement strain assessing oxidative and thermal stress sensitivity. Plates were photographed after 3 days. Images are representative of three biological replicates. (B) Polysome profiles of wild type (left) and the *eIF3d*Δ mutant (right) grown to mid-log at 30°C (unstressed) or treated with 1mM or 1.5 mM peroxide for 30 minutes (top) or shifted to 37°C for 30 or 60 minutes (bottom). Traces are representative of three biological replicates.

### Deletion of eIF3d leads to reduced eIF2α phosphorylation in response to stress

The sensitivity of the eIF3dΔ mutant to 3-AT and peroxide are traits shared in common with loss of *GCN2*, the sole eIF2α kinase encoded in the *C. neoformans* genome. This led us to explore Gcn2 activation in this mutant using phosphorylation of eIF2α as a readout. In response to escalating doses of hydrogen peroxide, the eIF3d mutant was found to be defective in the induction of eIF2α phosphorylation whereas the wild type KN99 cells exhibit a dose-dependent increase in eIF2α phosphorylation (Fig. 4A). The striking defect in eIF2α phosphorylation was surprising given the wild type translational repression seen by polysome profiling in the mutant. We then probed the same protein with an antibody to total eIF2α and small ribosomal subunit Rps6 to assess the overall abundance of each between wild type and *eIF3d*Δ and found that there was no obvious difference in total eIF2α between strains, and a modest reduction in RPS6 abundance in the eIF3dΔ mutant. This suggests that the residual eIF2α phosphorylation in the *eIF3d*Δ mutant is sufficient to achieve translational repression, or that eIF3d plays an opposing role to translational repression that is relieved in its absence.

**Figure 4:**
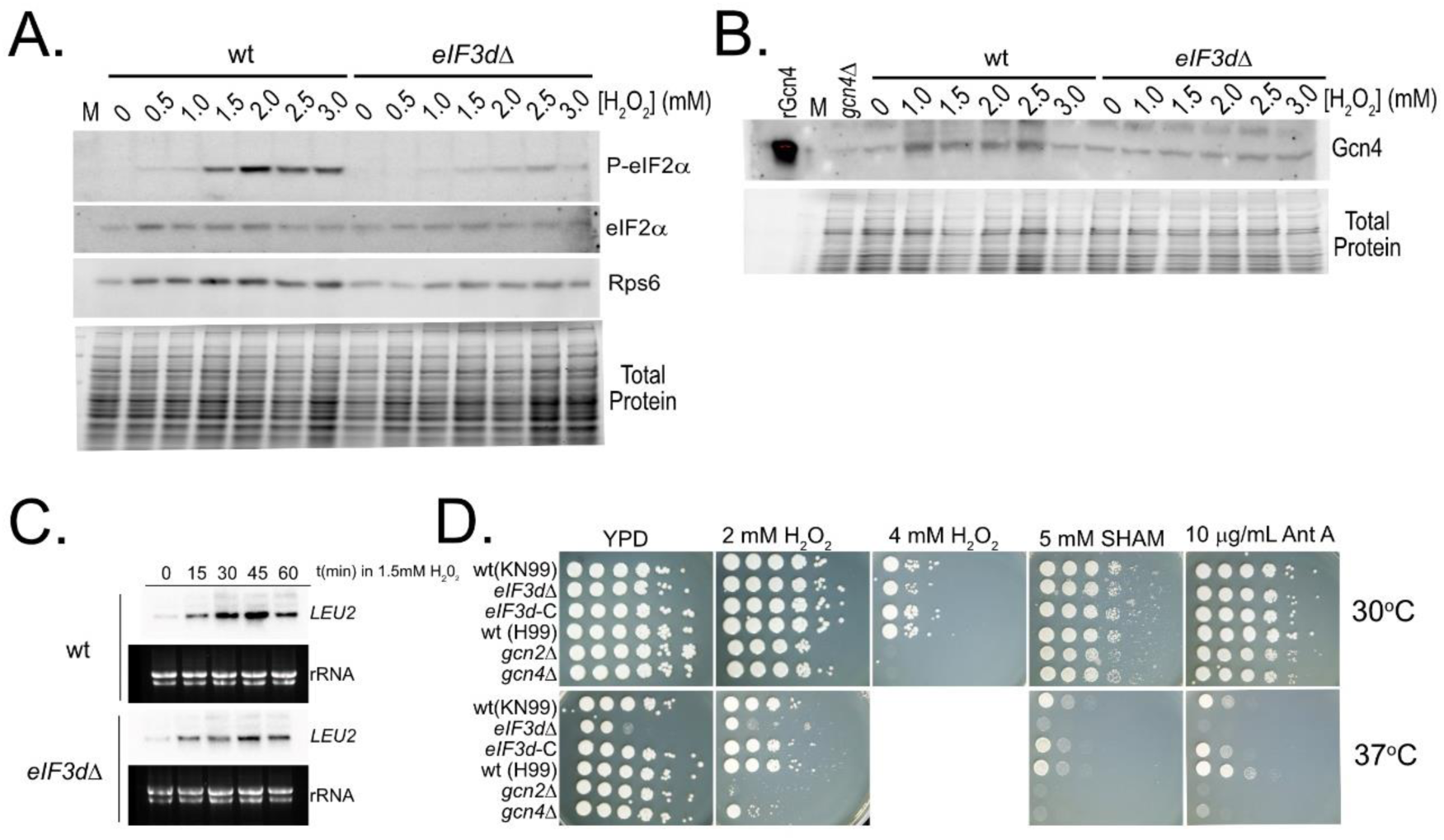
eIF3d is required for wild type Gcn2 activation and ISR induction. (A) western blot analysis of phosphorylated eIF2α, total eIF2α, and Rps6 in cultures of the wild type (left) and *eIF3d*Δ mutant (right) grown to mid-log at 30°C and treated with the indicated concentrations of peroxide for 30 minutes. (B) Western blot analysis assessing induction of the Gcn4 in the wild type (left) and *eIF3d*Δ mutant (right) grown to mid-log at 30°C and treated with the indicated concentration of peroxide for 30 minutes. (C) Northern blot analysis for the induction of the ISR target *LEU2* in the wild type (top) and *eiF3d*Δ mutant (bottom) in a time course following treatment with 1mM peroxide. (D) Spot plate analysis comparing the sensitivity of the *eIF3d*Δ mutant to the *gcn2*Δ and gcn4Δ mutants. Plates were photographed after 3 days of incubation. Respective isogenic wild type strains are included. All blot and plate images are representative of three biological replicates.

### ISR induction in the absence of eIF3d is intact but attenuated despite absence of a robust Gcn2 response

The result of Gcn2 activation is two-fold in that it represses cap-dependent translation by limiting 43 S preinitiation complex availability but also increases the stringency of start codon selection leading the production of Gcn4 via upstream open reading frame (uORF) bypass (23). Gcn4 is the transcription factor that drives induction of the integrated stress response (ISR) which we have reported to govern amino acid biosynthesis and redox balance in *C. neoformans* (4, 6). We assessed Gcn4 production by western blot in a similar peroxide dose response and found that Gcn4 protein was induced by peroxide in the wild type but was unchanged in the absence of eIF3 (Fig. 4B). Gcn4 drives transcription of ISR target genes, and so we assessed ISR induction in a time course following peroxide treatment by measuring expression of *LEU2*, a strongly induced gene involved in leucine biosynthesis. *LEU2* expression was induced strongly in the wild type in a time course following treatment with 1.5mM peroxide, and although there was induction in the *eIF3d*Δ mutant, it was to a lesser extent than in the wild type (Fig. 4C), consistent with a defect in ISR induction.

Given that the *eIF3d*Δ mutant represses translation as the wild type, but exhibits defects in ISR induction, we compared the stress phenotypes of the *eIF3d*Δ mutant to that of the *gcn2*Δ and *gcn4*Δ mutants. We found that the *eIF3d*Δ mutant exhibits an intermediate phenotype to that of *gcn2*Δ and *gcn4*Δ with respect to peroxide sensitivity, but the thermal stress sensitivity of *eIF3d*Δ appears independent of Gcn2-related defects. Interestingly, deletion of *GCN2* and *GCN4* result in a temperature-induced sensitivity to both SHAM and antimycin A which was also seen in the *eIF3d*Δ mutant (Fig. 2A and 4D), and suggests that both Gcn2 and eIF3 are connected to mitochondrial function under thermal stress. Future work is necessary to determine the specific regulon of each that impacts mitochondrial function.

### Loss of eIF3d enhances melanin and urease production but impairs virulence

We next set out to investigate a potential role for eIF3d in *C. neoformans* pathogenesis. Both melanin production and urease production are inducible virulence factors, with melanization requiring induction of *LAC1* in response to carbon starvation, and the urease activity being induced in the presence of urea (24). To our surprise, the absence of eIF3d increased melanization and urease production both at 30°C and 37°C (Fig. 5A and B). Melanization was not observed in the presence of 2% dextrose, demonstrating that the increased melanin production is not due to derepression of laccase expression under conditions of carbon catabolite repression. We next used the *G. mellonella* model of infection to investigate virulence (Fig. 5C), and found that although the wild type and complemented strains were lethal to the larvae within six days, no larvae infected with the *eIF3d*Δ mutant or saline died in the same time period.

**Figure 5.**
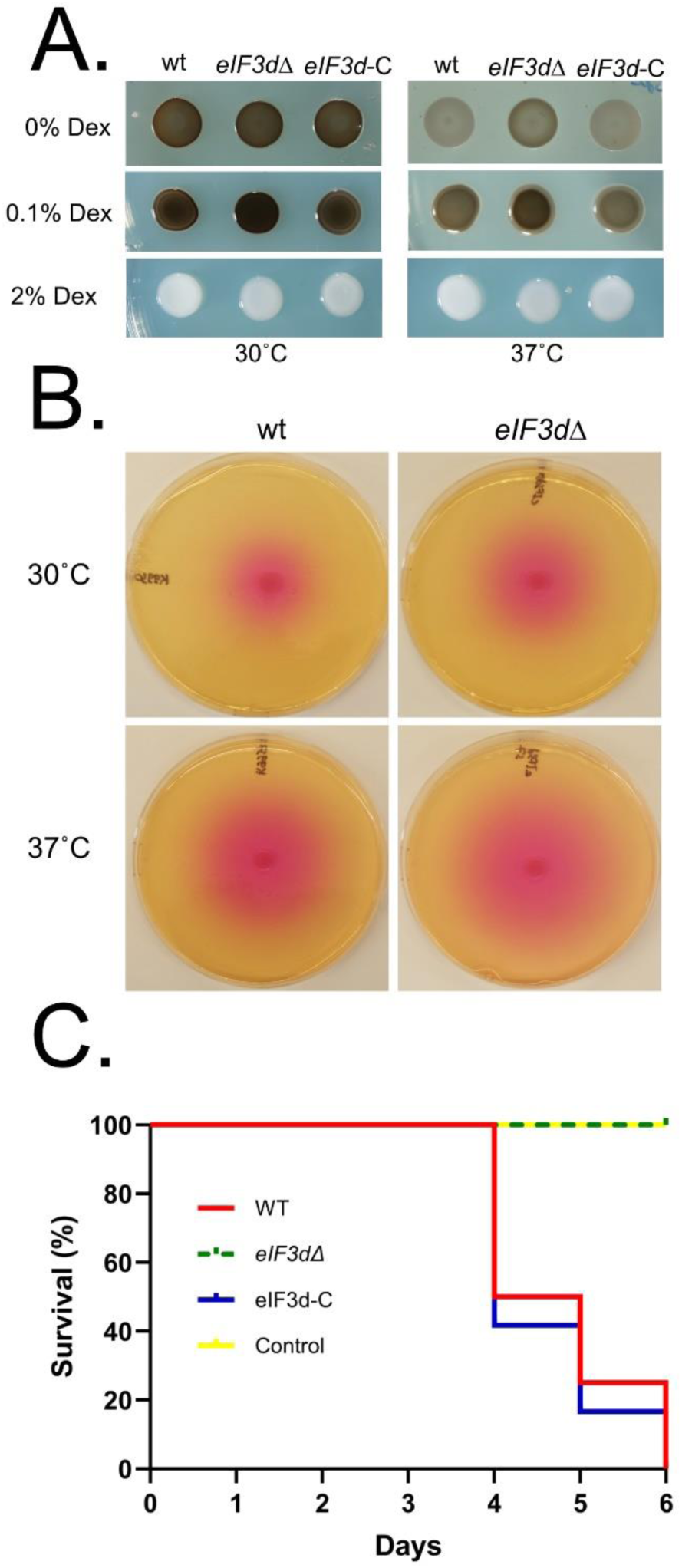
eIF3d regulates virulence factor production and virulence in an invertebrate model of cryptococcosis. (A) Melanin production assays using L-DOPA as substrate indicated glucose concentrations and temperatures. Plates were photographed after 24 hr. Images are representative of three biological replicates. (B) Urease activity assays on Christenson’s Agar incubated at indicated temperatures. Photographs were taken after 18 hours. Images are representative of three biological replicates. (C) Mortality curves from infection of *Galleria mellonella* (wax moth larvae) with wild type *C. neoformans* KN99, the *eIF3d*Δ mutant, or the complemented strain and incubated at 37°C (bottom) for six days. Curve is a representative of three biological replicates.

In sum, our data demonstrate that *C. neoformans* encodes a mammalian eIF3 complex, and that the non-essential subunits contribute to stress adaptation. Further investigation of eIF3d demonstrated that despite normal translational repression, the induction of the integrated stress response was defective. Further work is necessary to define the specific regulons of Gcn2/4 and eIF3d that contribute to oxidative stress and mitochondrial stress adaptation in the context of thermal stress.

## Discussion

Stress adaptation is a hallmark trait of pathogenic fungi, and the regulation of stress adaptation occurs at multiple levels along the path of gene expression, from the regulation of mRNA synthesis to the endpoint decisions of mRNA fate (4, 7, 25). One understudied area of stress adaptation in fungal pathogens is ribosome selectivity under stress. eIF3, the largest and most complex eukaryotic translation initiation factor, has functions that bridge mRNA binding and translation initiation that implicate it in ribosome selectivity (10, 26). Further, dissociable subunits may promote translation of specific mRNAs under stress conditions. For example, eIF3d in *S. pombe* is required for translation of mRNAs encoding metabolic enzymes and components of the electron transport chain (13).

We undertook a sequence-based inquiry of genomes of both model fungi and fungal pathogens, to determine the composition of eIF3. Of the fungi assessed, only *S. cerevisiae* and *N. glabratus* contained the minimal eIF3 with only six subunits. Given that *N. glabratus* is capable of causing disease in humans, it appears that eIF3 component composition is not universally predictive of pathogenic potential. *C. albicans* and *C. auris* each include a single MPN domain-containing protein, eIF3f, and its associating factor, eIF3m. The ascomycete molds, including *N. crassa* and *A. fumigatus* encode the full mammalian complement of eIF3 subunits, as do all of the basidiomycete fungi interrogated.

Taking advantage of the *C. neoformans* deletion collection, we identified deletion mutants in four of the eIF3 subunits, eIF3d, e, k and l. Interestingly, three of these are PCI-containing proteins that form the mammalian core, and excludes the two PCI proteins that are shared between mammals and *S. cerevisiae.* The remaining mutant was in eIF3d, which is a dissociable subunit that is thought to interact with the PCI core on the mRNA exit channel side of the ribosome, but this is based on additional density within the observed cryo-EM structure of eIF3 in complex with 40 S ribosome subunit (16). A published signature-tagged mutagenesis (STM) screen in *C. neoformans* identified eIF3l as important for lung infectivity with a significant depletion in the STM output, and thus was named *LIV12* (27).

Although all four mutants exhibited some degree of stress sensitivity, loss of eIF3d led to the most robust stress-sensitive phenotype that included sensitivity to oxidative stress, translational stress and thermal stress (28, 29). Our previous work has implicated the Gcn2 kinase in regulating the response to oxidative stress in *C. neoformans* (4). Work from mammalian cells has implicated eIF3d in regulating Gcn2 activation and integrated stress response induction after chronic ISR-inducing stress, which led us to investigate Gcn2 activation and ISR induction in the absence of eIF3d (29). We were surprised to find that translational repression in response to oxidative and thermal stress, as observed by polysome profiling, was completely intact in the absence of eIF3d. This suggests that there is sufficient signal to repress translation despite severely attenuated phosphorylation of eIF2α in the *eIF3d*Δ mutant. We did, however, observe a defect in ISR induction both at the level of Gcn4 production and induction of the ISR transcript *LEU2.* This is consistent with a role for eIF3 in regulating uORF bypass that has been demonstrated previously with ATF4 in mammalian cells (30).

Our previous work with Gcn2 in *C. neoformans* demonstrates that in the absence of additional stressors, the role of Gcn2 and the ISR in thermal stress is minimal (4, 6). Thus, the severe thermal stress sensitivity of eIF3d is likely independent of regulation of ISR induction. This suggests additional roles for eIF3d in *C. neoformans* that are related to thermal stress adaptation and pathogenesis. We explored two virulence factors that produced by *C. neoformans,* melanin and urease, and found the activity of these factors was increased in the absence of eIF3d. However, using the *G. mellonella* model of *Cryptococcal* infection, we found that the *eIF3dΔ* mutant was unable to cause mortality in the larvae incubated at 37°C. This is consistent with recently published phenome data that demonstrate a fitness defect of the *eIF3d*Δ mutant in mice (31). Interestingly, an eIF3d deletion mutant was unable to be generated in *N. crassa,* suggesting that eIF3d is essential in this ascomycete mold (14). Further work is necessary to determine the specific mRNAs in *C. neoformans* that rely on eIF3d for selective translation under stress to promote adaptation and pathogenesis.

## Acknowledgements

This work was supported by PHS Grant R01AI157459 to JCP. The Madhani deletion collection was supported by NIH R01AI100272 to Hiten Madhani, UCSF and obtained through the Fungal Genetics Stock Center at Kansas State University.

